# Astroglial deficiency for oligophrenin-1 contributes to intellectual disability

**DOI:** 10.1101/2025.01.09.632152

**Authors:** Noémie Cresto, Pascal Ezan, Laure-Elise Pillet, David Mazaud, Philippe Mailly, Alexis Bemelmans, Hamid Meziane, Mohammed Selloum, Tania Sorg, Nicola Marchi, Yann Hérault, Pierre Billuart, Nathalie Rouach

**Author notes:** Contributed equally (Pierre Billuart, Nathalie Rouach). Institute of Functional Genomics, University of Montpellier, CNRS, INSERM, Montpellier, France.

## Abstract

Neurodevelopmental disorders, including X-linked intellectual disability, autism spectrum disorder or schizophrenia, can result from the mutation of oligophrenin-1 (*Ophn1*), encoding a Rho-GTPase-activating protein. *Ophn1* regulates synaptic development and function, in part via cytoskeleton reorganization, and is expressed in both neurons and astrocytes. Despite the crucial role of astrocytes in synapse function, altered in neurodevelopmental disorders, and their *Ophn1* expression, the specific impact of astroglial *Ophn1* deficiency on synaptic transmission and behavior remains unknown. Here, we show that *Ophn1* deficiency postnatally in hippocampal astrocytes impairs synaptic transmission, short-term plasticity and spatial working memory in adults. This involves an adenosine A1 receptor-dependent presynaptic mechanism associated with astroglial morphological rearrangements resulting in increased astroglial synapse coverage. The structural, functional and behavioral alterations induced by astroglial *Ophn1* deficiency are rescued in adults by pharmacological inhibition of the RhoA/ROCK pathway. Our findings uncover an important role for astroglial *Ophn1* deficiency in synaptic and behavioral dysfunctions, pointing to a novel cellular therapeutic target for neurodevelopmental disorders.

## INTRODUCTION

Intellectual disability (ID) is a neurodevelopmental disorder that manifests before the age of 18, leading to impaired intellectual functioning (IQ < 70) and deficits in adaptive behavior, encompassing social, conceptual, and practical skills^1^. Affecting 1–2% of the global population, ID is frequently misdiagnosed, resulting in substantial healthcare costs and underscoring the urgent need for innovative therapeutic approaches^2^. The etiology of ID is multifaceted, involving both genetic and environmental factors.

In males, the second most prevalent form of genetic ID, after Down syndrome, arises from mutations in genes located on the X chromosome^3^. A significant proportion of these ID-related genes encode proteins critical for synaptic function, with many cases of ID, particularly X-linked ID (XLID), being associated with abnormalities in gene expression regulation (chromatinopathies) or synaptic dysfunctions (synaptopathies)^4–6^. The *Oligophrenin-1* gene (*Ophn1*) is implicated in a syndromic form of XLID accompanied by cerebellar hypoplasia, and missense mutations in *Ophn1* have also been linked to schizophrenia and autism spectrum disorders (ASD)^7–9^. This association, supported by a high score from the Simons Foundation Autism Research Initiative (SFARI), highlights *Ophn1* as a strong candidate gene. *Ophn1* is highly expressed at synapses and modulates the RhoA/ROCK/MLC2 signaling pathway through its RhoGAP domain, playing a pivotal role in cytoskeletal remodelling^10,11^. Recent findings have demonstrated that constitutive *Ophn1* deficiency during postnatal development leads to progressive deficits in synaptic transmission and plasticity, primarily due to presynaptic dysfunction^12^.

Although much research has focused on the involvement of neurons in ID, the intricate structure of the synapse-comprising pre- and postsynaptic elements surrounded by astroglial processes-raises questions about the role of glial cells, particularly astrocytes, in the pathology. Notably, genes associated with ID/ASD, historically considered neuronal, have recently been found to be also expressed in astrocytes (∼70% of ID-related genes)^13–16^, as shown in studies on *Ophn1* expression in human cells^17,18^ and to regulate their morphology *in vitro* and *in vivo*^17^. Indeed, O*phn1* deficient astrocytes isolated from constitutive *Ophn1* knockout (KO) mice, display an altered morphology resulting from the hyperactivation of the RhoA/ROCK/MLC2 pathway^17^. Astrocytes actively interact with neurons, influencing synapse formation, maturation, receptor trafficking, and synaptic activity by releasing neuroactive molecules that target pre- or postsynaptic elements^19,20^. Additionally, astrocytes regulate synaptic efficiency by controlling extracellular glutamate levels through astroglial transporters, whose activity depends on their proximity to perisynaptic processes^21–23^. Dysregulation of these astroglial processes may contribute to the synaptic abnormalities found in the *Ophn1* knockout mouse model of XLID and, more broadly, in ID and ASD.

In this study, using conditional, local and inducible *Ophn1* deficiency in adult hippocampal astrocytes, we reveal that astroglial Ophn*1* plays a crucial role in regulating hippocampal synaptic transmission, plasticity and spatial memory. This regulation is associated with changes in astrocyte morphology and synaptic coverage. Mechanistically, the RhoA/ROCK signaling pathway mediates the structural alterations of astrocytes and the functional deficits in synaptic transmission and memory induced by astroglial *Ophn1* deficiency.

## RESULTS

### Conditional, local and inducible *Ophn1* deficiency in adult hippocampal astrocytes

To investigate the effects of *Ophn1* deficiency in astrocytes *in vivo*, we generated an astroglial conditional, inducible, and local *Ophn1* knockdown mouse (KDastro) using local Cre/Lox mediated recombination via viral vectors (Fig. 1a, b). To this end, we expressed locally and specifically in hippocampal astrocytes either GFP-tagged Cre-recombinase (Cre-GFP) or GFP only, as a control, under the GFAP (glial fibrillary acidic protein) promoter by delivering adeno-associated virus serotype 2/9 (AAV2/9) in the hippocampus of P15 *Ophn1*^fl/y^ or WT mice, respectively (Fig. 1a). Three months after the viral injection, we confirmed in KDastro mice the inversion of the *Ophn1* flox sequences after Cre expression using PCR on genomic DNA from injected hippocampi and detected the recombined allele from the inverted locus only in the injected hemisphere (Fig. 1c). We also verified the astrocytic specificity of *Ophn1* deletion by co-immunolabeling of GFP and Cre-recombinase with astroglial (GFAP) or neuronal (NeuN) markers. Single cell analysis in *Ophn1*^fl/y^ mice injected in the hippocampus with AAV2/9 GFAP-Cre-GFP revealed that 86% of GFAP-positive cells were also positive for Cre-recombinase and GFP (Fig. 1d, e), whereas only 3% of NeuN positive cells expressed the Cre-recombinase (Fig. 1f, g), confirming specific viral targeting of astrocytes.

**Fig. 1.**
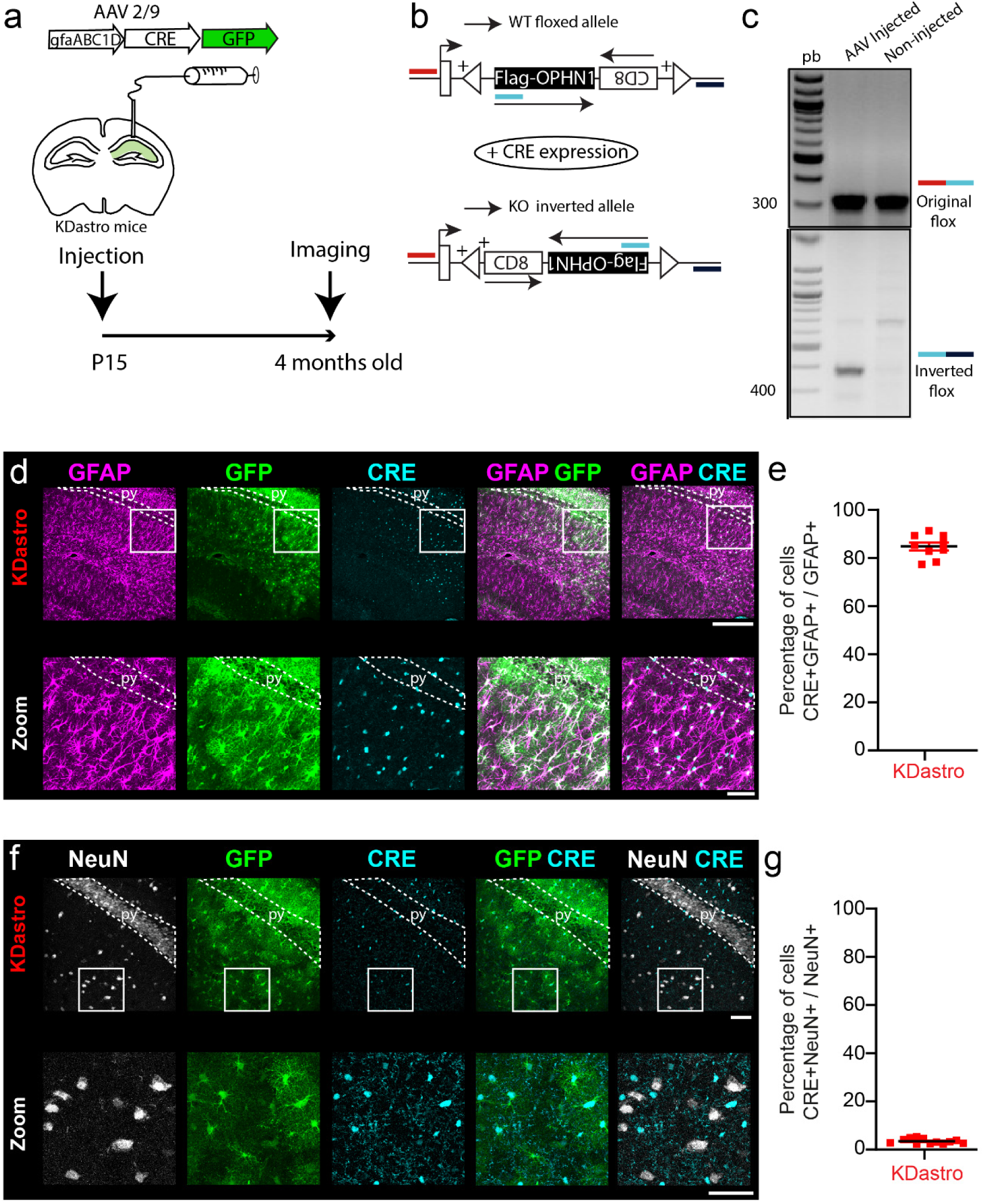
Generation of an astroglial conditional, inducible and local *Ophn1* knockdown mice (KDastro) **a,** Diagram of AAV2/9 vectors designed to express the Cre-recombinase and a GFP reporter under the GFAP promoter (gfaABC_1_D) and scheme of unilateral AAV2/9 microinjection into the mouse hippocampus. *Ophn1* floxed mice were injected at P15 and imaging was performed 3.5 months post-injection. **b,** The floxed allele of *Ophn1* consists of the *Ophn1* cDNA inserted in the *Ophn1* locus in the first intron. This cassette is flanked by inverted LoxP sites, which leads to the inversion of the cDNA upon Cre-recombinase and inactivation of *Ophn1* in Cre-expressing cells. **c,** PCR detection of the original flox allele (top panel in b) in both hemispheres (combination of red and blue primers) and the inverted flox allele (bottom panel in b) only in the AAV-injected side (combination of blue and purple primers). **d-e,** The expression of the GFP and Cre-recombinase was primarily detected in GFAP positive cells (n=9 ROI from 3 mice), **f-g,** but was barely detected in NeuN positive cells (n=12 ROI from 4 mice). Scale bars: (d) 500 µm; (zoom) 40 µm; (f) 40 µm; (zoom) 40 µm.

### Astroglial *Ophn1* deficiency imparis spatial working memory and synaptic transmission

Previous studies have demonstrated a role for *Ophn1* in spatial working memory, which is impaired in constitutive *Ophn1* KO mice. To investigate the contribution of astrocytes to this memory deficit, we tested the performance of KDastro mice, deficient for *Ophn1* selectively in hippocampal astrocytes, in the Y maze. We found in these mice that spatial working memory is abolished (Fig. 2a, b), as reflected by the percentage of spontaneous alternation in the Y-maze around chance level (50%), in contrast to control mice (WT*-GFP*). We also tested the selective contribution of neuronal *Ophn1* in spatial working memory by using the *CamKII-Ophn1^fl/y^* mice. We found no defect in spontaneous alternation in the Y maze in these mice compared to floxed mice without Cre (Supplemental Fig. 1, a), pointing to the selective role of astroglial *Ophn1* in spatial working memory.

**Fig. 2.**
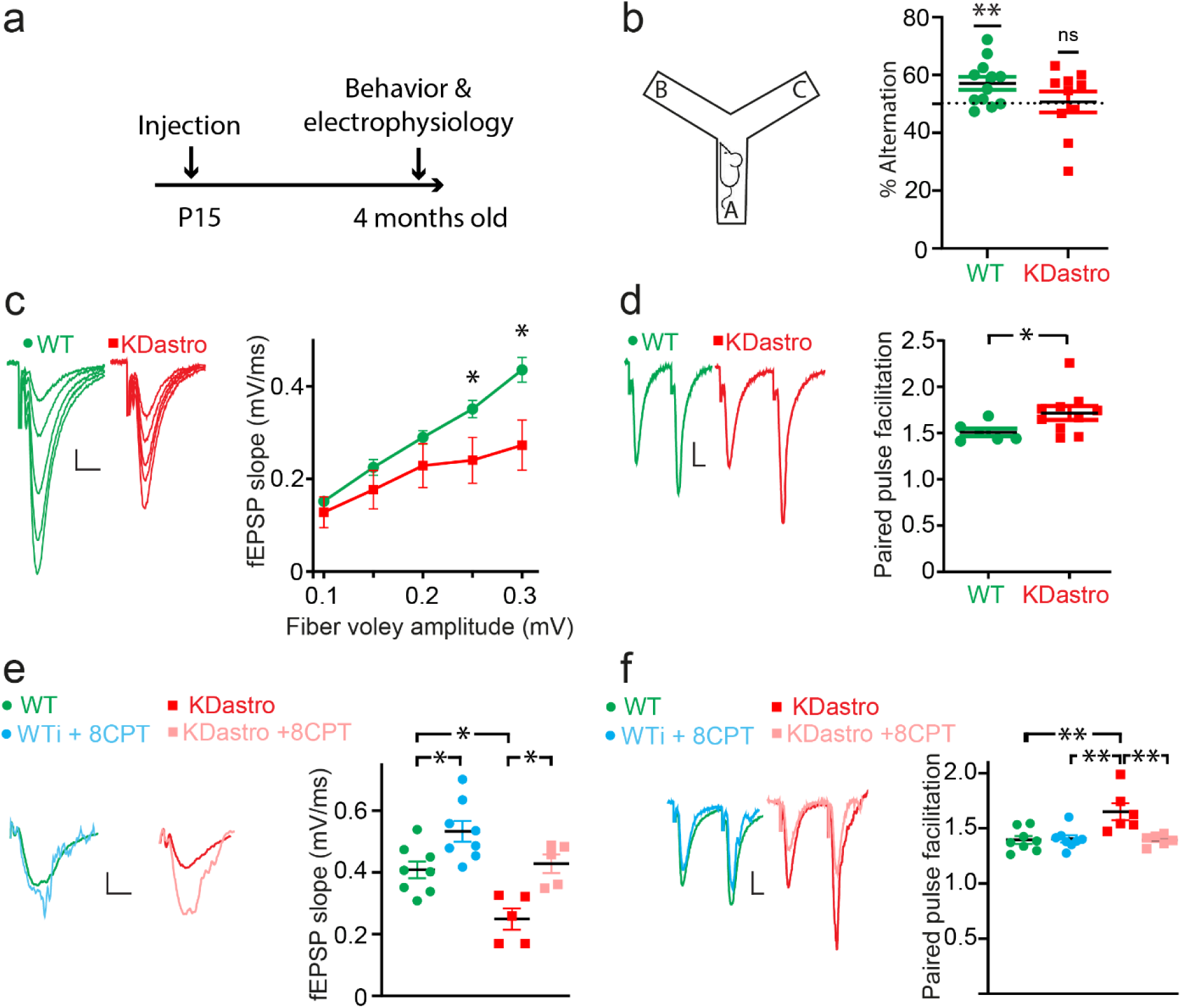
Astroglial *Ophn1* regulates synaptic transmission. **a,** Scheme illustrating the viral injection (AAV2/9-gfa-Cre-GFP) in P15 mice followed 3.5 months later by behavioral (Y-maze) and electrophysiological analysis (basal synaptic activity in the injected area). **b,** Spatial working memory was evaluated using the Y-maze in the spontaneous alternation configuration (control WT n=12 mice, KDastro n=10 mice, one sample t-test compared to 50 as hypothetical mean. Control WT p= 0.0089, KDastro p= 0.8525). **c,** Electrophysiological analysis with input/output curve showing that basal excitatory synaptic transmission is reduced in KDastro mice as compared to control WT mice (n=7 mice, KDastro n=8 mice, two-way ANOVA, Bonferroni post-hoc test, p=0.0135 at fiber volley amplitude=0.3 mV, scale bar: 0.2 mV; 20 ms). **d,** The paired pulse facilitation (PPF) is increased in KDastro mice, as compared to WT control mice (WT n= 6 mice, KDastro n=10 mice, Mann Whitney test, p= 0.0312, scale bar: 0.1 mV; 10 ms). **e-f,** Basal excitatory synaptic transmission and PPF was assessed before and after 8-cyclopentyl-theophylline (8-CPT) application, an A1R antagonist. **e,** Control WT n=8 mice; KDastro n=5 mice; One way ANOVA, ****: p<0.0001 and Bonferroni post-hoc test. control WT vs KDastro, p= 0.0148; WT vs WT + 8-CPT, p=0.0358; KDastro vs KDastro + 8-CPT, p=0.0136, scale bar: 0.2 mV; 20 ms). **f,** WT n=8 mice, KDastro n=6 mice, one-way ANOVA, **: p: 0.0012 and Bonferroni post-hoc test. WT vs KDastro, p=0.0028; KDastro vs KDastro + 8-CPT, p=0.005, scale bar: 0.1 mV; 10 ms).

We then investigated at the synaptic level the basis for this deficit. Constitutive *Ophn1* KO mice display impaired excitatory synaptic transmission due to presynaptic defect^12,24–28^. We thus assessed basal excitatory synaptic transmission at CA1-Schaffer collateral synapses by recording field excitatory postsynaptic potentials (fEPSP) in acute hippocampal slices from adult WT mice or *Ophn1^fl/y^* mice both transduced with Cre-GFP (*KDastro* mice) (Fig. 2, a). We found in adult KDastro mice a reduction in excitatory synaptic transmission by comparing the size of the presynaptic fiber volley (input) to the slope of the fEPSP (output) (Fig. 2c). To determine whether neurotransmitter release was decreased in adult KDastro mice, we recorded paired-pulse facilitation (PPF), a form of short-term plasticity sensitive to changes in release probability. PPF was increased in KDastro mice (Fig. 2d), pointing to a decrease in the probability of presynaptic release. It is noteworthy that PPF was unchanged at earlier developmental stage, i.e. in juvenile mice (Supplemental Fig. 2). Taken together, these results suggest that *Ophn1* deficiency in hippocampal astrocytes from adult mice replicates the alterations in excitatory synaptic transmission found in constitutive *Ophn1* KO mice^12^, underscoring a critical role for astroglial *Ophn1* at the presynapse.

We then investigated the molecular mechanism mediating the presynaptic regulation by astroglial *Ophn1*. Among the gliotransmitters negatively regulating the presynaptic function, ATP can be metabolized into adenosine, which activates presynaptic A1 receptors (A1R) to inhibit neurotransmitter release^29–32^. We thus tested the implication of this pathway in KDastro mice by determining the effect of the endogenous activation of A1R on excitatory synaptic transmission and release probability. To this end, we inhibited the adenosine A1R with 8-cyclopentyl-theophylline (8-CPT), and found in KDastro mice that this fully rescued to wildtype level excitatory synaptic transmission and release probability, as assessed with fEPSP and PPF recordings (Fig. 2, e, f). In contrast, inhibiting A1R in control mice (WT-GFP) had no effect on PPF, and only moderately increased fEPSP (∼+30%), compared to the massive increase found in KDastro mice (∼+72%) (Fig. 2e, f). Altogether, these data indicate that astroglial *Ophn1* decreases excitatory synaptic transmission via a presynaptic mechanism mediated by activation of adenosine A1 receptors.

### *Ophn1* controls astrocytes morphology and synapse coverage

*Ophn1*, through its RhoGAP domain, plays a crucial role in remodeling the cytoskeleton^11^. In addition, structural changes in astrocyte morphology and coverage of synapses can impact neurotransmission by controlling the levels of neuroactive molecules in the extracellular space ^21,22,33,34^. We thus tested whether *Ophn1* regulates the structural properties of astrocytes and of their interactions with synapses in adult mice, i.e. a stage at which *Ophn1* impacts synaptic transmission. To this end, we performed 3D reconstruction of hippocampal astrocytes expressing GFP, which revealed that *Ophn1*-deficient astrocytes display an increased volume, total process length and branching complexity compared to WT astrocytes (Fig. 3a-d). In addition, we found similar structural changes in astrocytes from constitutive *Ophn1* KO mice (Supplemental Fig. 3a). The increased volume and complexity of *Ophn1*-deficient astrocytes may result in a closer proximity of astroglial processes to synapses. To test this hypothesis, we performed 3D reconstructions of tripartite excitatory synapses using super-resolution STED imaging of astroglial processes around pre- and postsynaptic elements (Fig. 4a, b). The cumulative and distribution frequencies of the distance of astroglial process to synapses indicate a closer proximity of astrocytes to synapses in KDastro mice compared to control WT mice (Fig. 4c, d). Similarly, a closer proximity of astroglial processes to synapses was found in the constitutive *Ophn1* KO mice (Supplemental Fig. 4a, b).

**Fig. 3.**
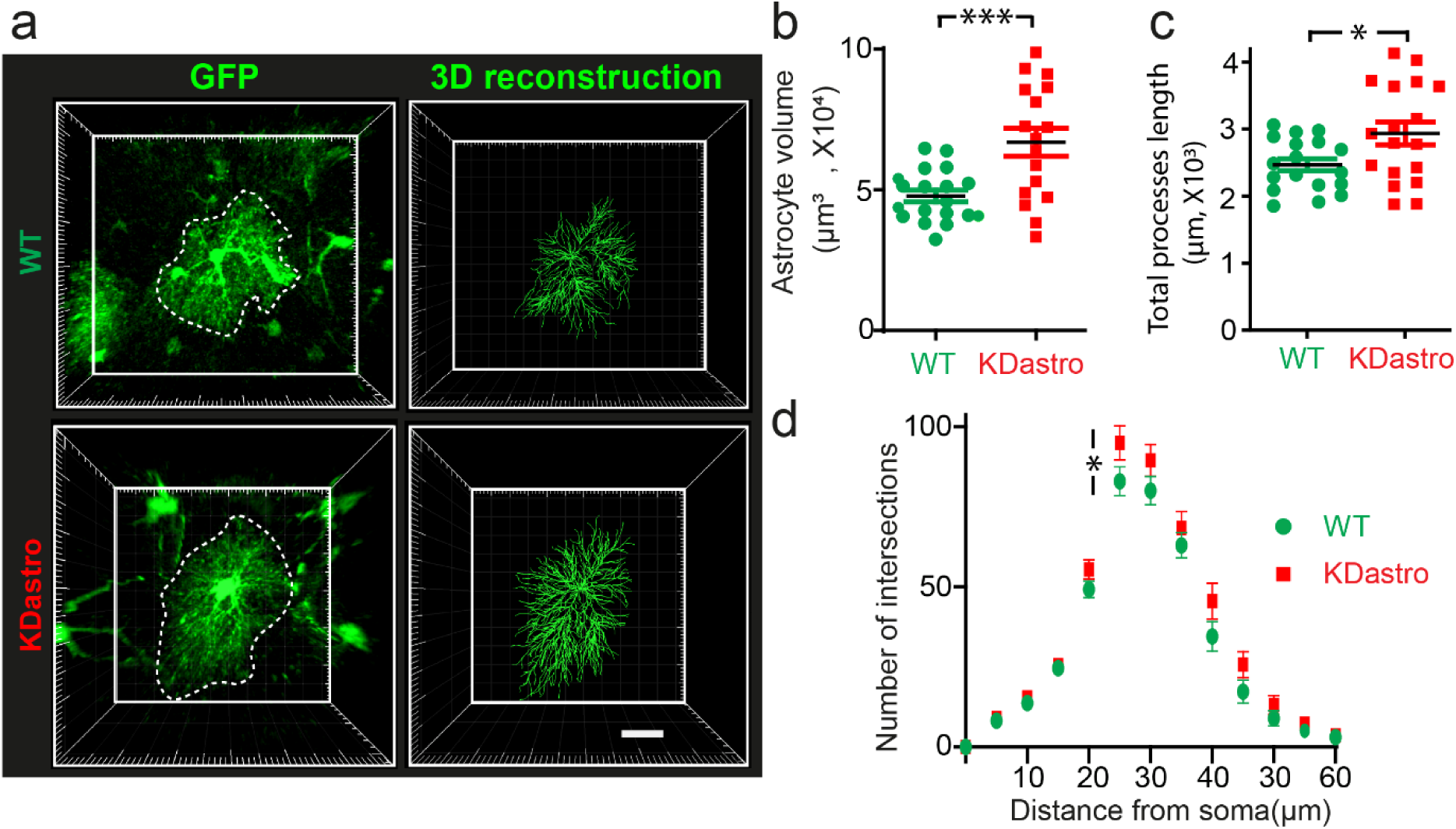
*Ophn1* controls astrocytes structural properties. **a,** Representative images of GFP-labeled astrocytes reconstructed in 3D. Scale bars: 10 µm. **b-d,** Quantifications of **b,** astrocyte volume (WT n=19 astrocytes, KDastro n=17 astrocytes, unpaired t test, p=0.0008), **c,** total process length (WT n=18 astrocytes, KDastro n=18 astrocytes, unpaired t-test, p= 0.0217) and **d,** process ramification using 3D-sholl analysis (WT n=19 astrocytes, KDastro n=18 astrocytes, two-way ANOVA, Bonferroni’s multiple comparison, p=0.0426).

**Fig. 4.**
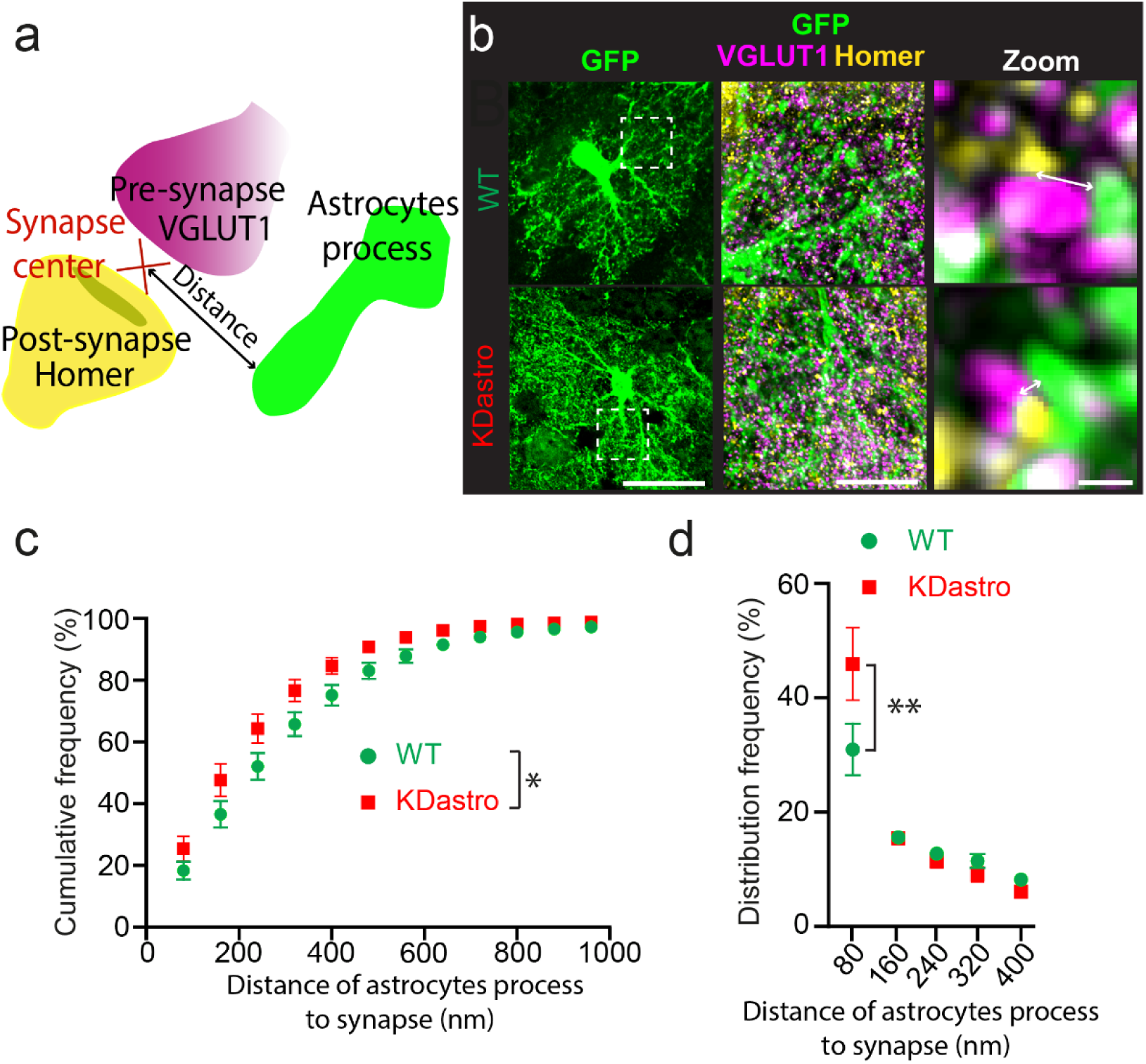
*Ophn1* controls astroglial synapse coverage. **a,** Scheme illustrating the coverage of glutamatergic synapses by astroglial processes and its quantification via the distance of astrocyte process to the center of the synapse. **b,** Representative images of GFP-labeled astrocytes (green) covering glutamatergic synapses labeled with VGLUT1 (presynaptic element, magenta) and homer (postsynaptic element, yellow). Scale bars: 30 µm (left), 10 µm (middle) and 1 µm (right). **c, d,** Quantification of astroglial coverage of excitatory synapses by the cumulative frequency and the distribution frequency of the distance of astroglial processes to synapses (**c**, WT n= 11 astrocytes, KDastro n= 15 astrocytes, Kolmogorov-Smirnov test, p= 0.0158, **d**, WT n= 11 astrocytes, KDastro n= 15 astrocytes, two-way ANOVA p=0.0112, Bonferroni’s multiple comparison p= 0.0015).

Altogether, these data indicate that astroglial *Ophn1* controls the geometry of astrocytes and their structural interactions with synapses.

### Activation of the RhoA/ROCK pathway mediates the effect of astroglial *Ophn1* deficiency at the structural, functional and behavioral levels

*Ophn1* regulates the RhoA/ROCK/MLC2 signaling pathway in astrocytes through its RhoGAP domain^35^. We thus here tested whether ROCK inhibition rescues the morphological, electrophysiological and behavioral changes observed in mice with conditional inactivation of *Ophn1* in astrocytes. To this end, we first evaluated astrocytic morphology in hippocampal slices acutely treated with Y27632. We found that the astrocytic volume was reduced in KDastro mice treated with Y27632, reaching a level similar to that found in WT hippocampal slices treated or not with Y27632 (Fig. 5a, b). Consistent with the decreased astrocytic volume induced by ROCK inhibition, the astroglial coverage of synapses was also reduced in KDastro mice to WT levels with or without Y27632 (Fig. 5c, d). It is noteworthy that no significant changes were observed when we compared the morphology and synapse coverage of WT and Y27632-treated WT hippocampal slices (Fig. 5b, d). Functionally, we found that ROCK inhibition fully rescued PPF to WT level in hippocampal slices from KDastro mice, assessed in the same recording before and after acute treatment with Y-27632 (Fig. 5e). Finally, at the behavioural level, ROCK inhibition also fully rescued spatial working memory, as assessed by analysis of spontaneous alternation in the Y-maze. The percentage of correct alternation in the Y-maze was indeed significantly above chance level (50%) in KDastro mice treated with Fasudil, in contrast to the untreated KDastro condition, reaching scores found in WT mice (Fig. 5f). Taken together, these findings indicate that targeting this RhoGTPase pathway rescues the structural, functional and behavioral effects of *Ophn1* deficiency in astrocytes.

**Fig. 5.**
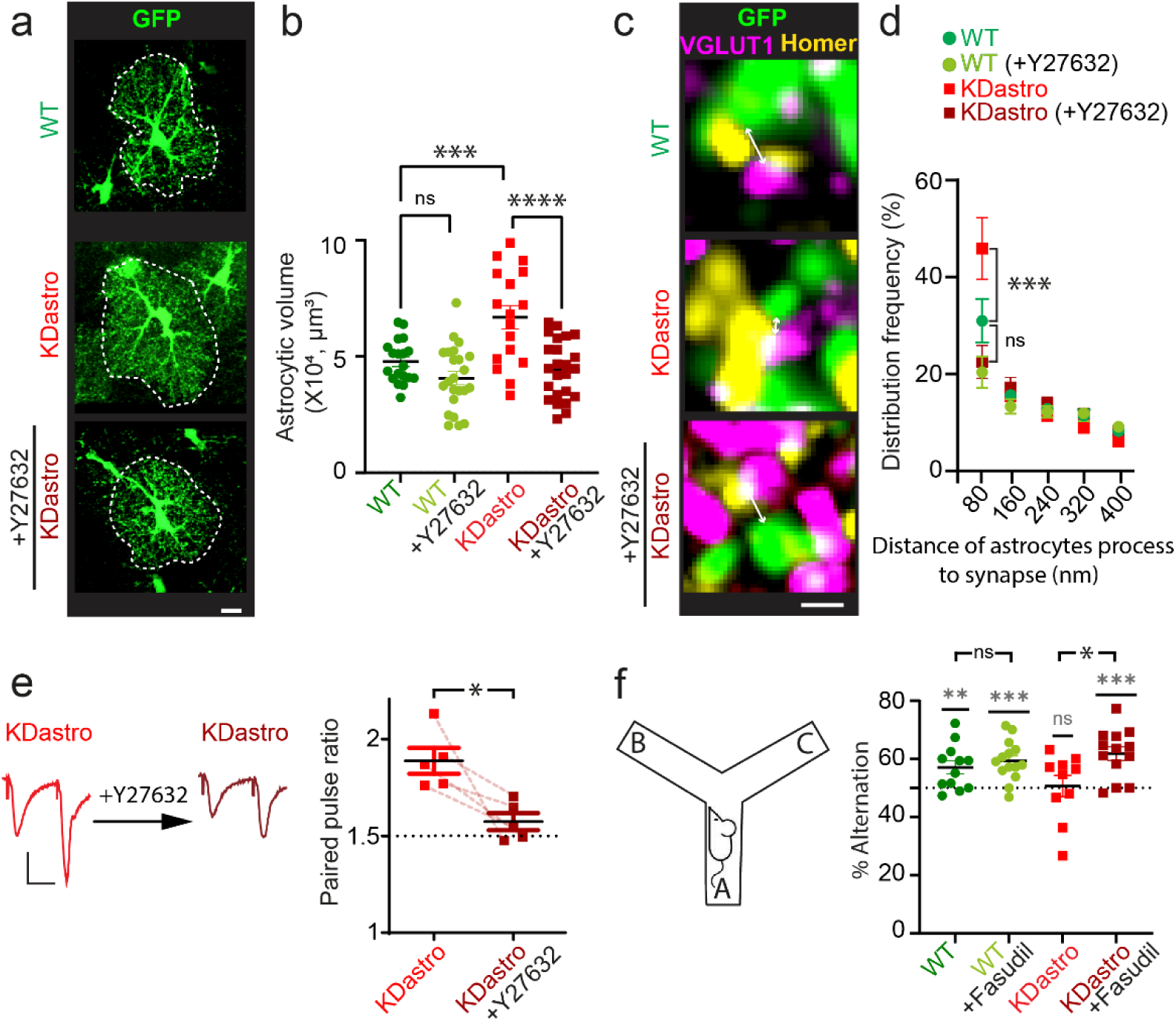
The structural and functional effects of astroglial *Ophn1* deficiency are mediated by activation of the RhoA/ROCK pathway. **a, b**, Quantification of the volume of GFP-positive astrocytes in acute slices treated or not with Y27632 (10 µM, 20 min) (WT, n= 19 astrocytes, KDastro, n= 17 astrocytes; WT + Y27632, n=24 astrocytes; KDastro + Y27632, n=24 astrocytes; ordinary two-way ANOVA test, p< 0.0001, Bonferroni’s multiple comparison, WT vs KDastro, p= 0.0008, KDastro + Y27632 vs KDastro, p<0.0001, WT vs WT + Y27632, p= 0.6801. ns: non-significant. Scale bars: 10 µm. **c, d,** Distribution frequency of the distance of astroglial processes to synapses in acute slices treated or not with Y27632 (WT, n=11; KDastro, n= 15; WT + Y27632, n= 10 astrocytes; KDastro + Y27632, n=12 astrocytes; Two-way ANOVA, p<0.0001, Bonferroni’s multiple comparison, WT vs KDastro, p=0.0001, KDastro + Y27632 vs KDastro, p<0.0001, KDastro vs WT + Y27632, p<0.0001. ns: non-significant. Scale bars: 1 µm. **e,** The PPF was measured in the same slice (KDastro) before and after Y27632 treatment (n=5). The dashed line represents WT level. Paired t-test, p=0.0293. Scale bar: 0.05 mV; 20 ms **f,** Spatial working memory was evaluated using the Y-maze in the spontaneous alternation configuration in mice treated or not with Fasudil in the drinking water (WT, n=12 mice; KDastro, n=14 mice; WT + Fasudil, n=10 mice; KDastro + Fasudil, n=13 mice). In light grey, One sample t-test, compared to 50 as hypothetical mean (WT, p= 0.0089; KDastro, p= 0.8525; WT + Fasudil, p=0.0002; KDastro + Fasudil, p= 0.0003). Ordinary two-way ANOVA test, p=0.0262, Bonferroni’s multiple comparison, WT vs KDastro, p= 0.5328, KDastro + Fasudil vs KDastro, p=0.0218, WT vs WT + Fasudil, p>0.9999. ns: non-significant.

## DISCUSSION

Here, we show that conditional inactivation of *Ophn1* in mouse hippocampal astrocytes alters basal synaptic transmission, short-term synaptic plasticity andspatial working memory, recapitulating the effects of the constitutive inactivation of *Ophn1* (*Ophn1-KO*). Interestingly, these effects are selective, as conditional inactivation of *Ophn1* in neurons from forebrains has no effect on spatial working memory nor on short-term synaptic plasticity. At the cellular level, *Ophn1* deficient astrocytes display changes in morphology consisting in an increased cellular volume, longer processes and higher ramification of middle branches. This translates at the synaptic level in a stronger coverage of synapses, with a shorter distance of the perisynaptic astrocyte processes to excitatory synapses. This closer synaptic proximity of astrocytes is associated with an increase in the activation of adenosine A1 receptors, resulting presynaptically in reduced neurotransmitter release. Finally, ROCK inhibitors treatment of mice deficient for astroglial *Ophn1* rescues the defects in astroglial morphology synaptic plasticity and behavior of mutant animals.

### Structural regulation of neuroglial interactions via the RhoA/ROCK in the *Ophn1* mouse model of intellectual deficiency

Here we report that the loss of *Ophn1* functions in astrocytes alters their morphology with increases in the volume, length, complexity of their processes and coverage of synapses. Since we found similar phenotypes in constitutive *Ophn1 KO*, this demonstrates a cell-autonomous function of *Ophn1* in the regulation of astrocyte morphology. All the morphological defects are rescued by the Y27632 ROCK inhibitor, indicating that deficiency for the RhoGAP *Ophn1* protein induces an activation of the RhoA/ROCK pathway in both, constitutive and astroglial conditional knockout mice. It has recently been reported that overexpression of constitutively activated variants of RhoA in hippocampal astrocytes reduces their volume fraction, selectively altering the peripheral and small astrocyte processes^36^. Indeed, the constitutively active form of RhoA selectively alters the peripheral and small astrocyte processes in vivo^36^. However, its effect on synapse coverage remains unknown. In contrast, we here report an overall increase in the volume of astrocytes, although we found that only astroglial middle branches, located at a distance >25µm from the soma, are more ramified. We also report at the synaptic level that deficiency for *Ophn1*, either constitutively (*Ophn1* Full KO) or selectively in astrocytes (KDastro), results in a closer proximity of the PAPs to the synapses.

These differential effects of RhoA activation on astrocyte structural properties may result from distinct patterns of RhoA signalling, according to the strength and spatial characteristics of its activation. Indeed, deficiency for *Ophn1*, a RhoGAP, leads to local activation of the RhoA pathway in some subcellular compartments enriched in F-actin, leaving the others unaffected ^11,37^. In contrast, overexpression of constitutively active RhoA strongly affects all compartments, and the resulting morphological changes are pleiotropic, leading to an overall decreased volume of astrocytes^36^.

In addition to the activation of the RhoA/ROCK pathway, we previously reported an increase in PKA activity in various brain areas of the constitutive *Ophn1 KO* model. This abolishes presynaptic long-term plasticity in the hippocampus and lateral amygdala, and increases signal noise in the medial prefrontal cortex (mPFC), which are associated with impairments in fear and working memory, respectively^28,38^. However, the cAMP levels are unaltered in the constitutive *Ophn1* KO, suggesting that the adenylate cyclase does not contribute to the increase in PKA activity^28^. Alternatively, ROCK and PKA share common substrates, and among them are GFAP and vimentin, which are key elements of intermediate filaments in astrocytes^39^. In addition to the canonical function of the RhoA/ROCK pathway in the regulation of the acto-myosin contraction, phosphorylation of key molecules of intermediate filaments may participate to the morphological changes observed in *Ophn1* deficient astrocytes, a phenotype which could be reversed by ROCK inhibitors.

### Functional regulations of neuroglial interactions and underlying mechanism in the Ophn1 mouse model of intellectual deficiency

We report here that either constitutive or local astroglial *Ophn1* deficiencies, decrease synaptic transmission and increase short-term presynaptic plasticity. Together these results suggest that a presynaptic function is altered in neurons secondary to *Ophn1* loss of function in astrocytes. Since PAPs in KDastro are closer to synapses than in controls, we tested the contribution of an altered gliotransmission associated to *Ophn1* deficiency. Amongst the gliotransmitters, ATP, which is released by astrocytes and metabolized into adenosine in the extracellular space, is known to activate A1 receptors at presynaptic sites. Activation of these Gi/o-coupled receptors at the presynapse inhibits adenylate cyclase and decreases cAMP levels, reducing transient calcium conductances and the release of glutamate by the ready releasable pool of vesicles^40^. Remarkably, we found that inhibiting A1 receptors in KDastro hippocampal slices, fully restored normal synaptic transmission and short-term plasticity. This indicates that A1 receptors are endogenously more activated upon *Ophn1* deletion in astrocytes. The closer proximity of the PAPs to the synapse may locally increases the ATP/adenosine levels in the extracellular space.

### Neuronal and astroglial controls of synaptic transmission and plasticity in the *Ophn1* model of intellectual deficiency

Studies on the constitutive *Ophn1* KO model in different brain regions (the hippocampus, lateral amygdala, olfactory bulbs, prefrontal cortex…) have shown that synaptic transmission and plasticity are altered in neurons, involving changes in presynaptic and/or postsynaptic functions^24,26–28,35,41–43^. More recently, it was demonstrated that the conditional inactivation of *Ophn1* in PV+ interneurons of the prefrontal cortex leads to a reduction in excitatory transmission, but without affecting the presynaptic release probability. This suggests a defect in the formation or stabilization of excitatory synapses in interneurons^44^. Here, we show that the conditional inactivation of *Ophn1* in hippocampal astrocytes reduces the presynaptic release probability due to an increase in A1 receptor signaling in the presynaptic compartment. Altogether, these results suggest that multiple mechanisms (vesicle secretion, recycling, and endocytosis, AMPA receptor endocytosis and stability…) involving different cell types (excitatory and inhibitory neurons, astrocytes) are at play in the pathophysiology of *Ophn1*-linked ID. However, in all cases, a common molecular mechanism involving the RhoA/ROCK pathway appears to emerge, as most of the observed phenotypes are corrected by ROCK inhibitors.

### Time window for *Ophn1* linked phenotypes and therapeutic intervention

We previously reported in vivo that astrocytes from constitutive *Ophn1* KO mouse have at P21 a similar morphology and complexity compared to control astrocytes^17^. We however here report in adults that constitutive or local conditional *Ophn1* deficiency in astrocytes both result in a larger volume of astrocytes. Additionally, we show that juvenile *Ophn1* KO mice had normal synaptic transmission and short-term plasticity, whereas in adult, they are decreased or increased, respectively^12^. Here we found similar results in *Ophn1* KDastro. It is noteworthy that all these neuronal changes coincide with the maturation of astrocytes, which peaks at one month of age^45^. This suggests that alteration of astrocyte development after birth participates to the neuronal deficit observed later. In addition, this temporal window may offer a good opportunity for postnatal therapeutic intervention in *Ophn1* linked ID.

## METHODS

### Ethics statement

All procedures on animals were performed according to the guidelines of European Community Council Directives of 01/01/2013 (2010/63/EU) and French regulations (Code Rural R214/87-130). Experimental procedures were approved by our local ethics committee (CEEA No. 059, Paris Centre et Sud) and registered with the French Research Ministry (APAFIS No. 2017112817353516).

### Animals

All applicable international, national, and/or institutional guidelines for the care and use of animals were followed. Adult mice were housed under a 12 h light/12 h dark circle, with ad libitum access to food and water. All efforts were made to minimize the number of used animals and their suffering, taking into consideration the “3Rs recommendation” (replacement, reduction, and refinement) for animal experimentation. Because the *Ophn1* gene is located on the X chromosome, only males *Ophn1* mice (*Ophn1*^tm1Bill/y^ *or Ophn1*^tm2Bill/y^) and wildtype (WT: *Ophn1*^+/y^) littermate were included in the study. *Ophn1*^tm1Bill/y^ and *Ophn1*^tm2Bill^ have been previously reported^41^; the first one *is a constitutive Knock-Out and the second* is conditional (*Ophn1*^fl/y^). Briefly, the floxed allele of *Ophn1* consists of the *Ophn1* cDNA from *Homo sapiens* inserted in the *Ophn1* locus that simultaneously disrupts the endogenous mouse gene and rescues it under *Ophn1* promoter. This rescue cassette is flanked by inverted and mutated LoxP sites, which leads to the non-reversible inversion of the cDNA upon Cre recombinase and inactivation of *Ophn1* in Cre expressing cells. This model has been previously validated *in vitro* using 4 hydroxy-tamoxifen to inactivate *Ophn1* in astrocyte from KDastro male mice (see fig1f and 1g in ^17^). In order to generate the neuronal model of *Ophn1* inactivation (KDneuron), CamKII-Cre mouse line was used to drive the loss of *Ophn1* function in neurons from forebrain^46^. Control male mice had the *Ophn1*^fl/y^ allele without the Cre recombinase.

### Genotyping

Various combinations of oligonucleotides allowed to differentiate the 3 different alleles of the *Ophn1* locus. Wildtype allele: intronUp4 (red)/mR4(blue) (150 base pairs (bp)) and Original flox allele: intronUp4/mR4 (300 bp) (mR4: tcc agc ggg gga tgc ccc at; intron Up4: acc tgc ctt cca aag gga gtg). Cre-mediated Inverted allele: mR4 (blue)/Intron Low1(purple) (400 bp). (Intron Low1: aca ggt ctt tgt tcc acg ctg aga cag).

### AVV production and injections

The AAV-GFAP-CRE-GFP construct consisted in a transgene composed, from 5′ to 3′, of CRE recombinase, the 2A peptide coding sequence and GFP cDNAs in a single open reading frame. The transgene was placed under the control of a GFAP-specific promoter in an AAV shuttle plasmid containing the inverted terminal repeats (ITRs) of AAV2^47,48^. The 2A signal allowed that GFP and CRE recombinase to be expressed as distinct polypeptides, thus preventing bias on CRE function and localization potentially induced by a fusion protein. The AAV-GFAP-GFP control vector encoded GFP only. Serotype 9 adeno-associated viruses (AAVs) particles were produced by transient co-transfection of HEK-293T cells, as previously described^49^. Viral titers were determined by quantitative PCR amplification of the ITR on DNase-resistant particles and expressed as vector genomes per ml (vg/ml). Juvenile mice were injected with AAV at P15 + 2 days. Animals were deeply anesthetized with a mixture of ketamine (95 mg/kg, Merial) and xylazine (10 mg/kg, Bayer) in 0.9% NaCl and placed in a stereotaxic frame, with constant body heat regulation. Injections were unilaterally performed in the right hippocampus at the following stereotaxic coordinates: AP: −1.85 mm; L: 1.6 mm to the bregma at a depth of −1.5 mm to the skull. We injected 1 μL of virus AAV-GFAP-GFP or AAV-GFAP-CRE-GFP at 1.10^10^ vg/mL, with a 29-gauge blunt-tipped needle linked to a 2-μL Hamilton syringe by a polyethylene catheter, at a rate of 0.1 μL/min, with an automatic pump. After the injection, the needle was left in place for 5 min before being slowly removed. The skin was glued, and mice recovery was checked for the next 24 h.

### Electrophysiology and acute slices treatment

#### Acute hippocampal slices preparation

Mice were sacrificed by cervical dislocation and decapitation. The hippocampi were rapidly isolated and sectioned at 4°C using a vibratome (VT1200S, Leica, Bannockburn, IL, USA) in an artificial cerebrospinal fluid (aCSF). For experiments on juvenile mice (25 to 30 days old), acute transverse hippocampal slices (400 μm) were prepared as previously described^21,50^. Slices were maintained at room temperature in a storage chamber in an aCSF containing (in mM) 119 NaCl, 2.5 KCl, 2.5 CaCl_2_, 1.3 MgSO_4_, 1 NaH_2_PO_4_, 26.2 NaHCO_3_, and 11 glucose, saturated with 95% O_2_ and 5% CO_2_, for at least 1 h prior to recording. For experiments on adult mice (3 to 4 months old), hippocampi were cut in a cold solution containing (in mM) 250 D-sucrose, 25 sodium bicarbonate (NaHCO_3_), 3 potassium chloride (KCl), 1 calcium chloride (CaCl_2_), 10 magnesium chloride (MgCl_2_), and 10 D-glucose, oxygenated with carbogen (95% oxygen (O_2_) and 5% carbon dioxide (CO_2_)). Slices were maintained at room temperature in a storage chamber for 30 min in the same solution and then transferred in normal aCSF, as previously described^51^.

#### Extracellular field potential recording

For field potential recordings, slices were transferred to a submerged recording chamber mounted on an Olympus BX51WI microscope equipped for infrared differential interference (IR-DIC) microscopy and perfused with aCSF containing picrotoxin (Sigma-Aldrich, P1675, 100 µM) at a rate of 1.5 mL/min by a peristaltic pump at 30-32°C. A cut was made between the CA3 and CA1 regions to prevent the propagation of epileptiform activity. Glass pipettes, pulled with a horizontal puller (Sutter Instrument, Novato, CA, USA) and filled with aCSF, were used to stimulate CA1 Schaffer collaterals and were placed at a distance of ∼200 µm from the area where evoked field excitatory postsynaptic potentials (fEPSP) were recorded. fEPSPs were recorded and filtered (low pass at 1 kHz) with an Axopatch 200 A amplifier (Axon Instruments, Union City, CA, USA), digitized at 10 kHz with an A/D converter (Digidata 1322 A, Axon Instruments), and stored and analyzed on a computer using Pclamp9 and Clampfit9 softwares (Molecular Devices, San Jose, CA, USA). Baseline evoked responses were monitored online over 10 min, and only slices with stable fEPSP amplitudes were included. Input–output (I-O) relations for fEPSPs were measured at the start of each experiment by applying a series of stimuli of increasing intensity to the Schaffer collaterals. Paired-pulse facilitation (PPF) of fEPSPs was evoked by delivery of two stimuli at an interval of 40 ms, and it was measured by dividing the peak amplitude of the second response to the one of the first response. For histology analysis, hippocampal slices were fixed in 2% paraformaldehyde (2% PFA diluted in PBS, Waltham, MA, USA).

### Histology, image acquisition and analysis

#### Immunofluorescence on PFA perfused brain

Mice were deeply anesthetized with Thiopental and perfused intracardially with PFA 2%. Brains were carefully dissected and post-fixed overnight with PFA 2% at 4°C. The next day, PFA was removed and the brains were cryoprotected overnight in phosphate buffer saline (PBS) containing 30% sucrose. Brains were sliced with a freezing microtome (30 µm). Slices were stored in a preservative solution containing (300 mL of glycerol, 300mL of ethylene glycol, 300 mL of TRIS-buffered saline and 100 mL of H_2_O). For immunofluorescence, slices were blocked for 1h at room temperature with phosphate-buffered saline (PBS, Sigma-Aldrich, Saint-Louis, MO, USA) containing 0.25% Triton X100 (Sigma-Aldrich, Saint-Louis, MO, USA) and 0.002% gelatin (Sigma-Aldrich, Saint-Louis, MO, USA), incubated 24h at 4°C with primary antibodies in the blocking solution (see antibody table for reference), washed three times with PBS, incubated overnight with the appropriate secondary antibodies in the blocking solution, and then washed three times with PBS before mounting with fluoromount (Invitrogen).

| antibody | specie | Dilution | reference |
| --- | --- | --- | --- |
| GFAP | Rabbit | 1:500 | Sigma G9269 |
| CRE | Mouse | 1:500 | MAB3120 |
| GFP | Chicken | 1:500 | Ab13970 |
| NeuN | Mouse | 1:500 | Merck MAB377 |
| Vglut1 | Mouse | 1:200 | Synaptic Systems,<br>#135511 |
| Homer1 | Rabbit | 1:200 | Synaptic Sytems,<br>#160003 |
| Goat anti rabbit Alexa<br>594 | Rabbit | 1:200 | Jackson<br>ImmunoResearch, #<br>111-586-045 |
| goat anti-Mouse<br>Alexa647 | Mouse | 1:200 | Life Technologies,<br>#A21235 |
| goat anti-mouse star<br>red | Mouse | 1/200 | Abberior #1001-500UG |
| goat anti-rabbit 594 | Rabbit | 1/200 | Invitrogen #A32740 |

#### Immunofluorescence on acute hippocampal slices

Mice were sacrificed by cervical dislocation and decapitation. The hippocampi were rapidly isolated and sectioned at 4°C using a vibratome (model VT1200S, Leica, Bannockburn, IL, USA) in an artificial cerebrospinal fluid (aCSF), after a recovery time, slices were fixed in 4% PFA during one night and stored in PBS. Slices (400 µm) were blocked for 1 h at room temperature with phosphate-buffered saline (PBS, Sigma-Aldrich, Saint-Louis, MO, USA) containing 1% Triton X100 (Sigma-Aldrich, Saint-Louis, MO, USA) and 0.2% gelatin (Sigma-Aldrich, Saint-Louis, MO, USA), incubated 48 h at 4°C with primary antibodies in the blocking solution, washed three times with PBS, incubated overnight with secondary antibodies in the blocking solution, and then washed three times with PBS before mounting with Abberior mounting medium (Abberior Instruments GmbH, Göttingen, Germany). To study astroglial morphology and synaptic coverage, astrocytes were labeled using a GFP antibody (1/500, green fluorescent protein, Abcam, Chicken, ab13970) and excitatory synapses were labeled using the presynaptic marker VGlut1 (in magenta, 1/200 Synaptic Systems, Göttingen, Germany, #135511) and the postsynaptic marker Homer-1 (in yellow, 1/200, Synaptic Systems, Göttingen, Germany, #160003). Appropriate secondary antibodies were then used (goat anti-mouse star red (1/200, Abberior, Göttingen, Germany, #1001-500UG), goat anti-rabbit 594 (1/200, Invitrogen, Waltham, MA, USA, #A32740).

#### Image Acquisition

For the analysis of the synapses coverage, images were taken using a super-resolution custom upright stimulated emission depletion (STED) microscope (Abberior Instruments GmbH, Göttingen, Germany). The microscope is based on a Scientifica microscope body (Slice Scope, Scientifica, Uckfield, UK) equipped with an Olympus 100X/1.4NA ULSAPO objective lens. It comprises a scanner design featuring four mirrors (Quad Scanner, Abberior Instruments GmbH, Göttingen, Germany), with 488 nm, 561 nm, and 640 nm excitation lasers available (Abberior Instruments, pulsed at 40/80 MHz). A laser at 775 nm (MPB-C, pulsed at 40/80 MHz) is used to generate STED beams. The conventional laser excitation and STED laser beams are superimposed using a beam-splitter (HC BS R785 lambda/10 PV flat, AHF Analysetechnik, Tübingen, Germany). Common excitation power with pulsed excitation ranges from 10–20 µW with STED power intensities of up to 200 mW in the focal plane. Nine regions of interest of about 400 to 700 µm^3^ were imaged in CA1. For sholl analysis and volume measurement, Z-stack images were acquired at a confocal microscope with a 40X magnification (Leica SP5 inverted confocal). All images used for analysis were taken with the same confocal settings (pinhole, laser intensity, digital gain, and digital offset).

#### Image Analysis

For the analysis of the synapse coverage, confocal images were deconvolved using Huygens software and combined with the STED images in one file for analysis. The analysis was performed in ImageJ with an in-house developed plugin. In brief, maxima intensities were identified in the STED images and then compared to the deconvolved confocal images to remove false-positive punctae. When two punctae from pre-(VGlut1) and postsynaptic markers (Homer-1) were within 200 nm of each other, a synapse was assigned as a pixel in between the two punctae and the distance between the synapse and the closest astrocytic processes was calculated. For 3D-sholl analysis and volume measurement, isolated astrocytes were selected based on their GFP staining. Then images were loaded on IMARIS software (Version 10, Oxford Instruments) to perform 3D-sholl analysis and volume measurement using convex hull calculation method that superimposed on the filament tracing. We used the filament reconstruction using the following custom settings: detect new starting points: largest diameter 7.00 μm, seed points 0.300 μm; remove seed points around starting points: diameter of sphere regions: 15 μm. Seed points were corrected for (either placed in or removed from the center of the somata) manually if the IMARIS algorithm placed them incorrectly. All surface and filament parameters were exported into separate Excel files and used for data analysis.

### Behavior

#### Spatial working memory

The apparatus used to test spatial working memory was a Y-maze made of plexiglas with three identical arms (40×9×16 cm) placed at 120° from each other. Each arm had walls with specific motifs allowing to distinguish it from the others. Each mouse was placed at the end of one of the three arms, and allowed to explore freely the apparatus for 5 min, with the experimenter out of the animal’s sight. Alternations were automatically measured as successive entries into each of the three arms as on overlapping triplet sets (i.e. ABC, BCA …). The percentage of spontaneous alternation was calculated as an index of working memory performance. Fasudil hydrochloride or HA1077 (F4660, LC laboratories, Boston, MA, USA) was given orally ad libitum in drinking water (100 mg/kg/d) seven weeks before behavioural experiments.

### Drugs

8-cyclopentyl-theophylline (8-CPT, Tocris, 6137, 1µM), trans-4-[(1R)-1-Aminoethyl]-N-4 pyridinylcyclohexanecarboxamide dihydrochloride (Y27632, Sigma-Aldrich, Y0503, 10µM).

### Statistical Analysis

All data are expressed as the mean ± *SEM*. Prior to statistical comparison, normality tests and variance analysis (Shapiro–Wilk test) were performed, and the appropriate parametric or nonparametric statistical test was used. Statistical significance for within-group comparisons was determined by two-way analysis of variance (ANOVA) followed by Bonferroni’s post hoc test, whereas Mann–Whitney or unpaired *t*-tests were used for between-group comparisons (*p* values are reported in the figure legends). Statistical analysis was performed using Prism 9.3.1 (Graphpad Software, CA, USA).

## Data availability

The data generated in this study are available in the main text and the Supplementary Information file. Source data are provided with this paper.

## Acknowledgments.

The authors thank all the members of the Neuroglial Interactions in Cerebral Physiology and Pathologies laboratory for scientific exchanges as well as the viral vector facility of MIRCen and the imaging and animal facilities of College de France for excellent technical assistance.

## Author Contributions

Conceptualization: N.C., P.B., N.R.; Formal analysis: N.C., L.E.P., P.M., H.M., M.S., T.S, Y.H., P.B., N.R.; Funding acquisition: P.B., Y.H., N.M., and N.R.; Investigation: N.C., L.E.P., H.M., M.S., T.S, P.B., N.R.; Resources: N.M.; Supervision: P.B., N.R.; Project administration: P.B. and N.R.; Writing original draft: N.C., P.B., N.R.; Writing – review & editing: all authors.

