## Supplementary Information for "Astroglial deficiency for oligophrenin-1 contributes to intellectual disability"

**This PDF file includes:**

Supplementary Figures 1-4

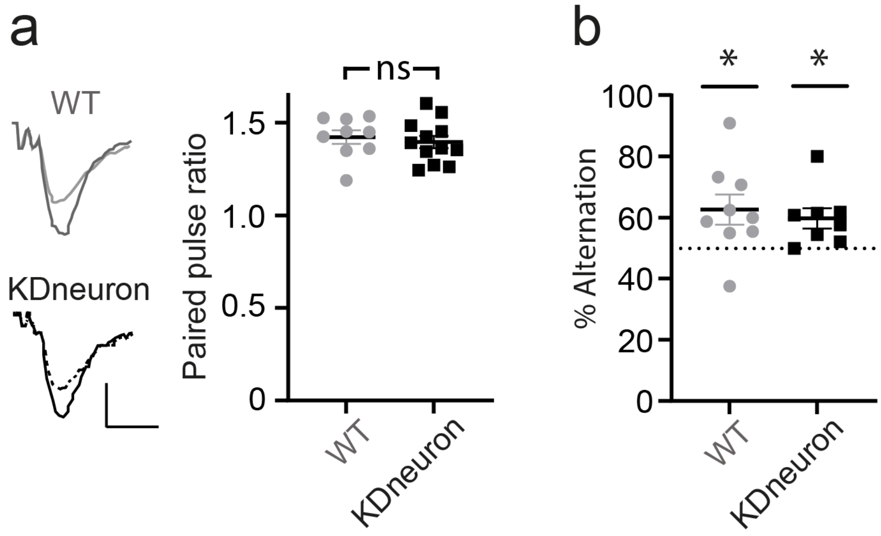


**Supplementary Fig. 1. Presynaptic function and spatial working memory are not altered in the Ophn1 conditional KO (KDneuron) mice driven by CamKII-cre. a,** The PPF was measured in slices from WT and KDneuron mice (WT, n= 9; KDneuron, n= 12; unpaired t-test, p=0.5939; *: p<0.01, ns: non-significant. **b,** The spatial working memory was evaluated using the Y-maze in the spontaneous alternation configuration (WT, n=9 mice, KDneuron, n=8 mice, one sample t-test, compared to 50 as hypothetical mean (WT, p=0.0324; KDneuron, p=0.0200, scale bar: 0.1 mV; 10 ms).

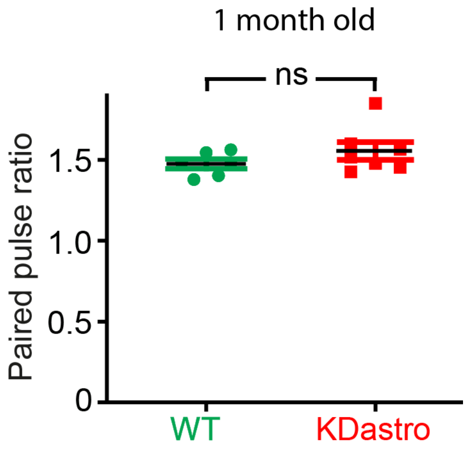


**Supplementary Fig. 2. Unaltered paired-pulse facilitation in juvenile knockdown mice for astroglial *Ophn1*.** The paired-pulse facilitation (PPF) was unchanged in KDastro mice, as compared to WT mice at 1 month-old (WT, n=6; KDastro, n= 7; unpaired t-test, p=0.2404; ns: non-significant).

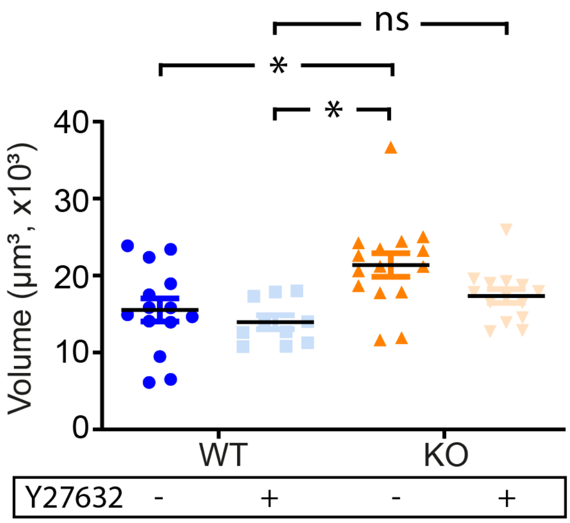


**Supplementary Fig. 3. The volume of astrocytes is increased via the RhoA/ROCK pathway in constitutive *Ophn1* knockout mice.** Quantification of the astrocyte volume in slices from WT and constitutive *Ophn1* knockout micetreated or not with Y27362, showing that deficiency for *Ophn1* increases the volume of astrocytes, which can be rescued to WT level with Y27632 (WT, n=14, KO, n=15, WT + Y27632, n=10, KO + Y27632, n=14; One-way ANOVA, p= 0.0016, Bonferroni's multiple comparison test, WT vs KO, p= 0.0114; WT + Y27632 vs KO + Y27632, p=0.5516 *: $p<0.01$).

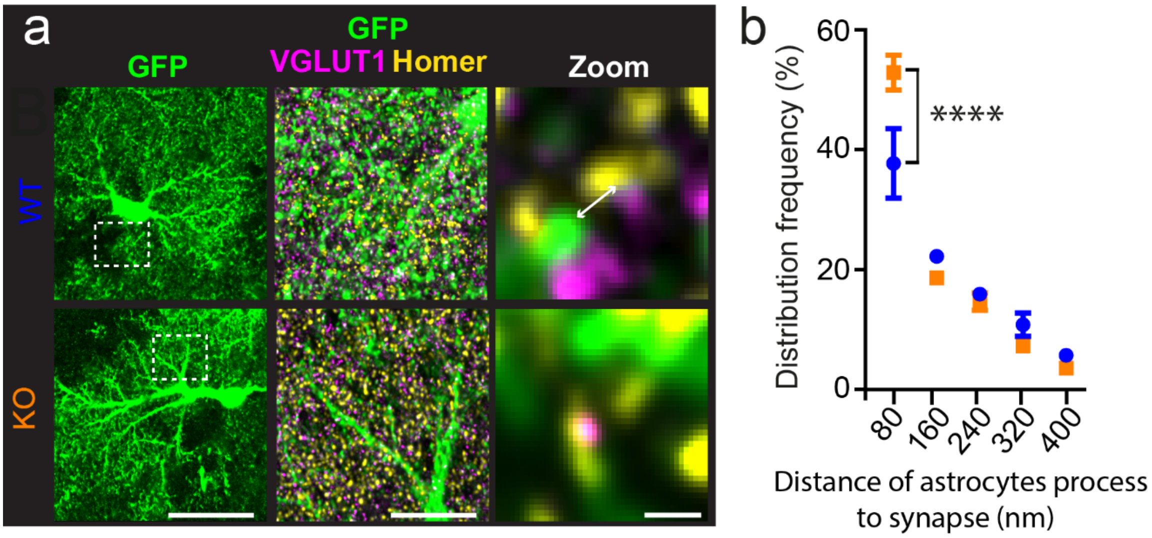


**Supplementary Fig. 4. *Ophn1* controls astroglial synapse coverage in constitutive *Ophn1* knockout mice. a,** AAV coding for GFP under the GFAP promoter was used to label the entire morphology of the astrocytes (GFP, green) in hippocampal slices. The excitatory presynaptic marker VGLUT1 (in magenta) and the postsynaptic marker Homer (in yellow) were used to label synapses. **b,** The distribution frequency was increased in the KO condition, suggesting that astrocytes processes are closer to synapses, as compared to the WT condition. Scale bars: 10 µm and 1 µm for the zoom. WT, n=4, KO, n=5. Two-way ANOVA, p< 0.003, Bonferroni's multiple comparison test, ****: p<0.0001.
